# The roles of vision and antennal mechanoreception in hawkmoth flight control

**DOI:** 10.1101/222448

**Authors:** Ajinkya Dahake, Anna Stöckl, James J. Foster, Sanjay P. Sane, Almut Kelber

## Abstract

Flying animals need constant sensory feedback about their body position and orientation for flight control. The visual system provides essential but slow feedback. In contrast, mechanosensory channels can provide feedback at much shorter timescales. How the contributions from these two senses are integrated remains an open question in most insect groups. In Diptera, fast mechanosensory feedback is provided by organs called halteres, and is crucial for the control of rapid flight manoeuvres, while vision controls manoeuvres in lower temporal frequency bands. Here we have investigated the visual-mechanosensory integration in an insect which lacks halteres: the hawkmoth *Macroglossum stellatarum*. They represent a large group of insects that use Johnston’s organs in their antennae to provide mechanosensory feedback on perturbations in body position. High-speed videos of freely-flying hawkmoths hovering at stationary or oscillating artificial flowers showed that positional fidelity during flight was reduced in flagella ablated animals, but was recovered after flagella re-attachment. Our experiments show that antennal mechanosensory feedback specifically mediates fast flight manoeuvres, but not slow ones. Differences in the latency of visual feedback (in different light intensities) affected all antennal conditions equally, suggesting there was no compensatory interaction between antennal and visual feedback under the tested conditions. These results establish the importance of antennal mechanosensors in providing rapid mechanosensory feedback for finer control of flight manoeuvres, acting in parallel to visual feedback.

## Introduction

The impressive aerobatic manoeuvres of insects provide an insightful model for the neural control of flight [Frye & Dickinson 2001, Fuller et al. 2014]. Insect flight requires constant sensory feedback, both on the position of the body relative to the environment, as well as on perturbations to body position. Visual feedback provides key information about flight parameters including ground speed, distance to obstacles and targets, and aerial displacements (for a review see [Srinivasan et al. 1999]). However, visual estimation of self-motion [Fuller et al. 2014, Hung et al. 2013] is limited by its temporal resolution and substantial latency to flight muscle activation [Sherman & Dickinson 2004, Suver et al. 2016]. This may often be too slow to control very fast aerial manoeuvres, which require rapid sensory feedback before perturbations become uncontrollably large and thus energetically costly to the animals [Bender & Dickinson 2006].

Avoiding the temporal limitations set by the visual system, insects use mechanosensors to sense their own motion, as these can transduce perturbations on much faster time scales [Yarger & Fox 2016]. The halteres of Dipteran insects are a classic example of gyroscopic function in active flight. Halteres are club-shaped mechanosensory structures that were evolutionarily derived from the hind-wings. They vibrate at the wing beat frequency and can sense rotations in any axis [Fraenkel & Pringle 1938, Nalbach 1994, Pringle 1948]. Halteres, however, are a special feature of only Dipteran (and Strepsipteran) insects [Pix et al. 1993]. How do flying insects from other orders, which also require fast feedback for stable flight, control flight manoeuvres without halteres? This question is especially interesting in insects active in dim light, as the visual systems of many insects trade off temporal acuity for sensitivity [Warrant 1999, Warrant 2017], thus rendering visually-based flight control even less reliable and increasing the need for mechanosensory feedback control.

Sphingids are a group of flying insects that are able to fly over a wide range of light intensities, due to their superposition compound eyes and additional neural adaptations [O’Carroll et al. 1996, O’Carroll et al. 1997, Stöckl et al. 2017, Theobald et al. 2010]. The effect of light intensity on their visual flight control has been quantified recently [Sponberg et al. 2015, Stöckl et al. 2017]. Moreover, the crepuscular hawkmoth *Manduca sexta* has been shown to use information provided by antennal mechanosensors, which may function similar to Dipteran halteres [Sane et al. 2007]. The mechanosensory Johnston’s organs, present at the pedicel-flagellar joint of the antennae, are stimulated by deflections of the antennal flagellum, are sensitive to a wide range of frequencies [Dieudonné et al. 2014], which far exceed the temporal response range of the visual system [Stöckl et al. 2017, Theobald et al. 2010]. After ablation of their flagella, the Johnston’s organs of *M. sexta* no longer receive relevant information, causing flight instability in these moths, whereas re-attachment of the flagella significantly improves their flight performance (Sane et al., 2007). Impaired flight performance following flagellar ablation was also observed in other Lepidopteran species, such as the tortoise-shell butterfly *Aglais urticae* [Gewecke & Niehaus 1981, Niehaus 1981] and the diurnal swallowtail moth *Urania fulgens* [Sane et al. 2010]. Although the above studies underscored the importance of antennal mechanosensors for natural flight, the severe behavioural impairment caused by flagellar ablation meant that the precise contributions of antennal mechanosensors to flight control remained an open question, as did their integration with the visual sense.

To address these questions, we therefore chose another model, the diurnal hawkmoth *Macroglossum stellatarum*, which retains both the motivation and ability to fly after flagella ablation. *M. stellatarum* feed from flowers while hovering in front of them, and previous studies have underscored the importance of visual feedback on their flower tracking behaviour [Farina et al. 1995, Farina et al. 1994, Kern 1998, Stöckl et al. 2017]. Using this model, we were able to test the role of antennal mechanosensors for the control of stationary hovering flight, as well as for flight manoeuvres at controlled temporal frequencies, focussing on the integration of visual and mechanosensory information (Fig. 1A). Here, we show that in *M. stellatarum*, antennal mechanosensors play a key role in the control of hovering flight, specifically in the control of fast flight manoeuvres (rapid turns). Furthermore, we show that visual and antennal mechanosensory feedback operate in different frequency bands, with no sign of compensatory interaction.

**Fig. 1.**
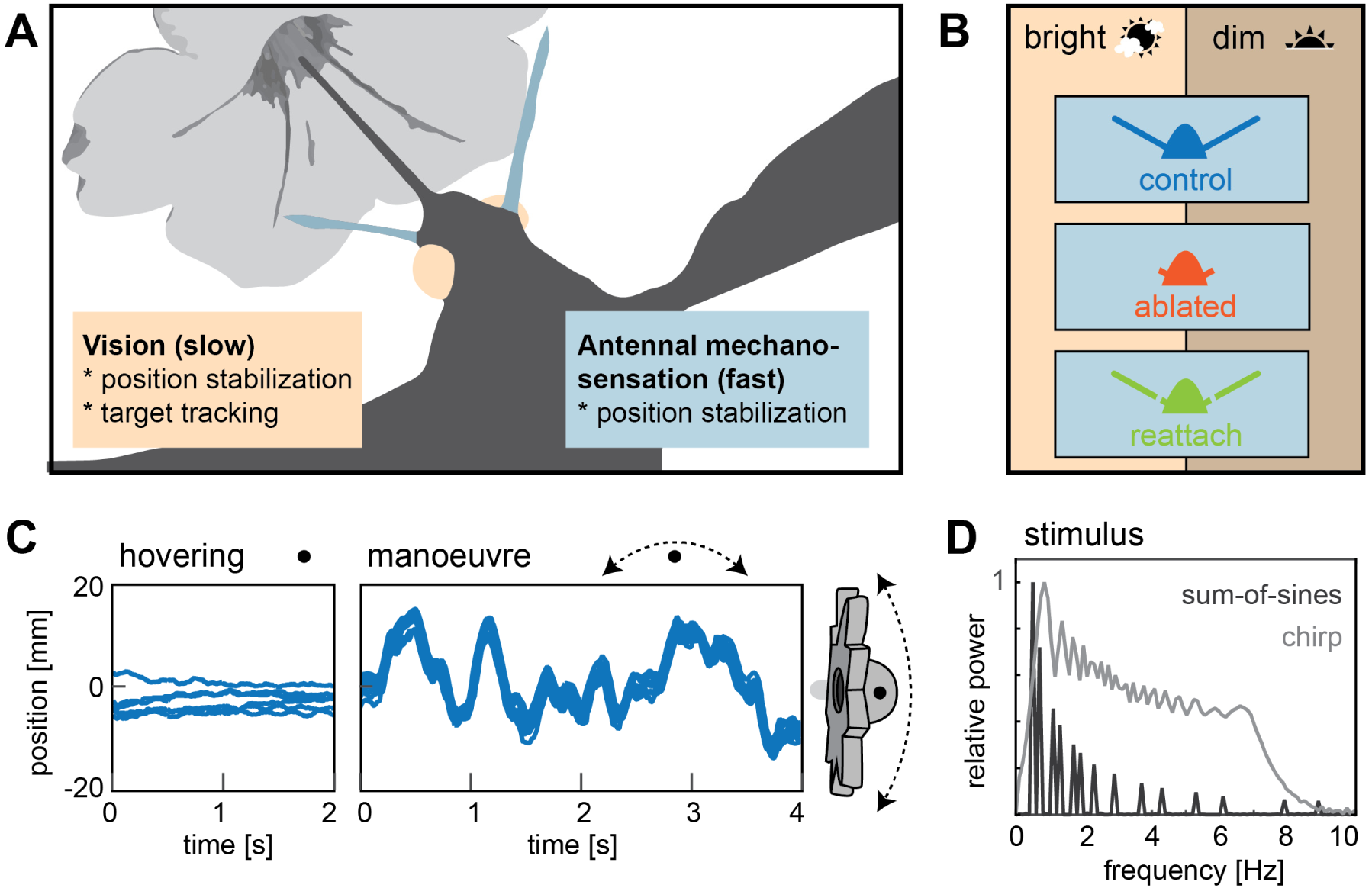
Flight control in hawkmoths requires vision and mechanosensation. **A**. Flight control in insects requires sensory feedback about perturbations in body position. The visual system supplies such feedback, but with comparably long response latencies. In addition, insects use mechanosensory systems to control their position in the air, which provide rapid feedback and thus are crucial for fast flight manoeuvres. Here, we investigated the role of antennal mechanosensation and vision on flight control in the hummingbird hawkmoth *Macroglossum stellatarum*. **B**. In order to quantify the effects of antennal mechanosensation in free flight, we subjected each hawkmoth to three treatments: intact antennae (*control*, blue), ablated flagella (*ablated*, red) and re-attached flagella (*reattach*, green). To quantify the role of vision, we tested these three antennal treatments in two different light intensities (*bright*: 3000lux, corresponding to partially overcast daylight and *dim*: 30 lux, corresponding to sunset intensities). **C**. All conditions were tested in free hovering flight at artificial flowers, which were either retained stationary (*hovering*) or moved at different temporal frequencies (*manœuvre*). **D**. We used a stimulus composed of a *sum-of-sines* to sample distinct frequencies with similar velocities (amplitude adjusted accordingly), as well as a stimulus ramping up in frequency, while retaining similar amplitude (*chirp*). See also Fig. S1.1

## Results

### Antennectomised hawkmoths performed less stable hovering at a stationary flower

To quantify the role of antennal mechanosensory feedback in free flight, we trained hawkmoths (*Macroglossum stellatarum*) in a flight cage to approach and hover in front of an artificial flower with sugar solution provided in a nectary at its centre (see Methods). Individual moths were tested in three antennal conditions: with intact antennae (*control*, blue, Figs 1B and S1.1), with ablated flagella (*ablated*, red) and with re-attached flagella (*reattach*, green). Rates of flower tracking decreased significantly with flagella ablation, but returned to control levels following reattachment (Table 1, S1). Yet, depending on light levels, a substantial proportion of individuals with ablated flagella still approached and fed from the flower (60% in bright and 36% in dim light) making it possible to study the combined roles of vision and antennal mechanosensory feedback on flight control in more detail.

**Table 1.** Proportion of moths performing specific behaviours across antennal conditions and light intensities. Proportion of trials in which animals performed the following behaviours: *no flight, flight* (but no tracking of the flower), *tracking*. This dataset is based on the animals participating in the moving flower experiments. Of the total numbers of moths, 27 control, 22 flagella ablated, and 14 re-attached moths were tested in both light intensities. Some moths were tested multiple times to collect the necessary tracking data, and thus have contributed multiple trials to this dataset. Statistical comparisons were performed using multinomial regression including the identity of individual moths as a random factor, to model the rates of one of the three behaviours as a function of antennal condition and lighting (without interaction terms). Statistical significance is indicated by: * p<0.05, ** p<0.01, *** p<0.001. For more details, see Table S1.

|  |  | control | ablated | reattach |
| --- | --- | --- | --- | --- |
| bright | No flight | 0.03 | 0.11 | 0.05 |
|  | Flight | 0.15 | 0.29 | 0.1 |
|  | Tracking | 0.82 | 0.60 *** | 0.85 |
|  | **Total** | **38** | **42** | **20** |
| dim *** | No flight | 0.03 | 0.34 | 0.08 |
|  | Flight | 0.09 | 0.30 | 0.08 |
|  | Tracking | 0.88 | 0.36 *** | 0.84 |
|  | **Total** | **35** | **66** | **25** |

When approaching the flower, moths with ablated flagella had distinctly longer and more tortuous flight trajectories than moths in the control and flagella re-attached conditions (Fig. S2.1). To further quantify flight performance, we focused on the hovering flight of the hawkmoth in front of a stationary or moving flower, where its body position could be closely monitored (Fig. 1C). Their target position was clearly defined by the position of the flower on which they fed. For the stationary flower experiment, we analysed the hovering flight of 6 animals, which performed in all three antennal conditions and both light intensities.

When hovering in front of the stationary flower, ablated moths jittered around their target position to a greater degree than moths in the other two antennal conditions, as evident from their thorax position over time (blue line, Fig. 2A). We quantified the amplitude of these thoracic movements across a range of frequencies from 0.5 to 50 Hz. In all three antennal conditions, the amplitude of thoracic jitter decreased with increasing frequencies (i.e. the animals performed smaller movements of higher frequencies) (Fig. 2B; Fig. S2.2A). At 3000 lux, the thoracic jitter of ablated moths was significantly larger than that of the other two antennal conditions between 0.7 and 5 Hz, and between 8 and 11 Hz (Fig. 2B, Table S2.1), whereas the difference between control and re-attached condition was not significant. Thus, re-attaching the flagellum restored flight performance close to the control state.

**Fig. 2.**
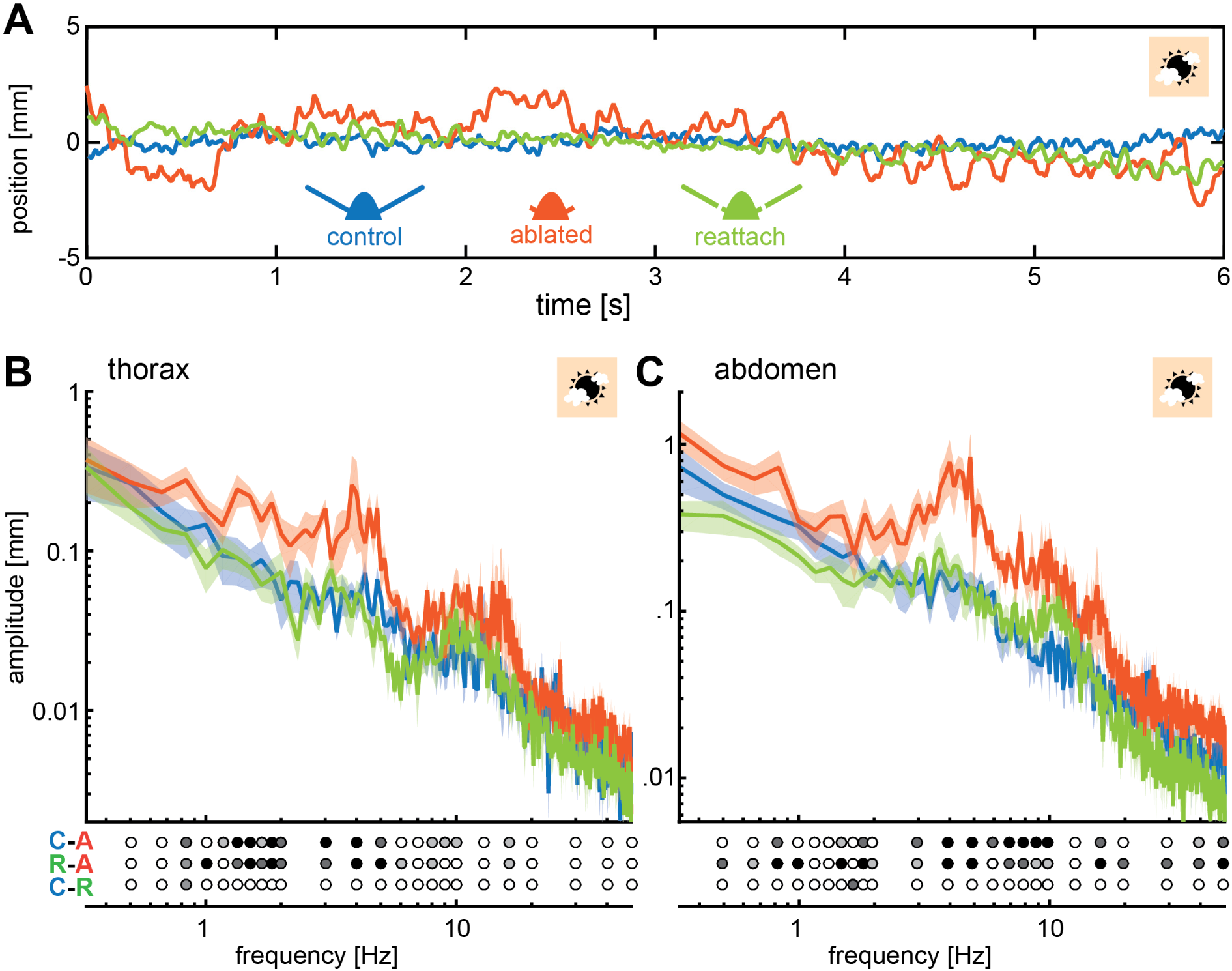
Flagella ablated hawkmoths showed greater thorax and abdomen movement during hovering flight at a stationary flower. **A**. When hawkmoths hovered in front of a stationary flower at 3000 lux, it was notable that flagella ablated moths jittered around their target position with larger amplitudes than moths of the other two antennal conditions, as quantified by the position of their thorax. The nectary is centered at 0 mm in this graph. **B**. The thorax of moths with ablated flagella jittered with significantly higher amplitudes than the other two antennal conditions at frequencies between 1 and 5 Hz. There was no significant difference between control and re-attached moths. Post-hoc tests were performed as part of a general linear model including antennal treatment and frequency (binned to the logarithmic scale) as factors. Statistical significance is indicated below the plot as: black p<0.001, dark grey: p<0.01, grey p<0.05, white p>0.05, see Table S2.1. **C**. Because hawkmoths are known to use abdomen movements to correct their position in the air, we also quantified the position of the abdomen in the three antennal treatments, showing a similar trend to the thorax: the flagella ablated moths exhibited significantly larger abdomen jitter in the frequency range between 0.5 and 10 Hz than the other two treatments (Statistical significance is indicated as in **B**, see Table S2.2). See also Fig. S2.1, S2.2

Because many insects, including hawkmoths, use abdominal movements for aerial stabilization during flight [Camhi 1970, Dyhr et al. 2013, Hinterwirth et al. 2012], we also quantified the movement of the abdomen over the same frequency range. Across all three antennal conditions, the abdominal and thoracic movements revealed similar trends; in flagella ablated moths, their magnitude was significantly larger than in control moths or flagella re-attached moths, over the entire range of frequencies tested (Fig. 2C, Table S2.2). Unlike thoracic jitter, abdominal jitter of moths with re-attached flagella differed significantly from control moths at only one frequency (1.66 Hz).

Since hovering is a dynamically unstable flight mode [Liang & Sun 2013, Wu & Sun 2012], hovering animals need constant sensory feedback to maintain a fixed position [Cowan et al. 2014]. Both the visual system and antennal mechanosensory systems could provide sensory feedback to correct for deviations from the target position. Because flagella ablated moths showed larger positional jitter, especially at higher frequencies, we conclude that antennal mechanosensory feedback is required for the control of hovering flight. Without antennal input, the feedback about deviations from the target position, likely supplied by the visual system and therefore slower, causes moths to drift further from their target position before a corrective manoeuvre can be initiated. This in turn results in greater thoracic and abdominal jitter.

### Flagella ablation reduces flower tracking performance at high frequencies in hawkmoths

After comparing the stationary hovering performance between antennal conditions, we examined the differences in performance at specific temporal frequencies (henceforth simply referred to as frequency). While the hawkmoths were feeding from the nectary, we moved the artificial flower laterally to elicit flight manoeuvres of controlled frequencies and amplitudes while the moths were tracking the flower (Fig. 1C). To probe the moth’s manoeuvrability at different frequencies, we used two movement patterns as stimuli: one pattern was generated from a sum of sine waves ranging from 0.5 to 8.9 Hz, which decreased in amplitude with increasing frequency, thus keeping a constant velocity (“sum-of-sines”, Fig. 3A). The second movement pattern had a constant amplitude, while its frequency increased over time from 0 to 7.3 Hz over time (“chirp”, Fig. 1D & 3C). We analysed the flight performance of 12 moths, which tracked both stimuli in all antennal conditions and two light intensities. Flight performance was analysed using a metric that evaluates their accuracy in tracking both the amplitude (Fig. S3.1D-F) and the phase (Fig. S3.1G-I) of the flower movement, termed tracking error [Roth et al. 2011, Sponberg et al. 2015].

**Fig. 3.**
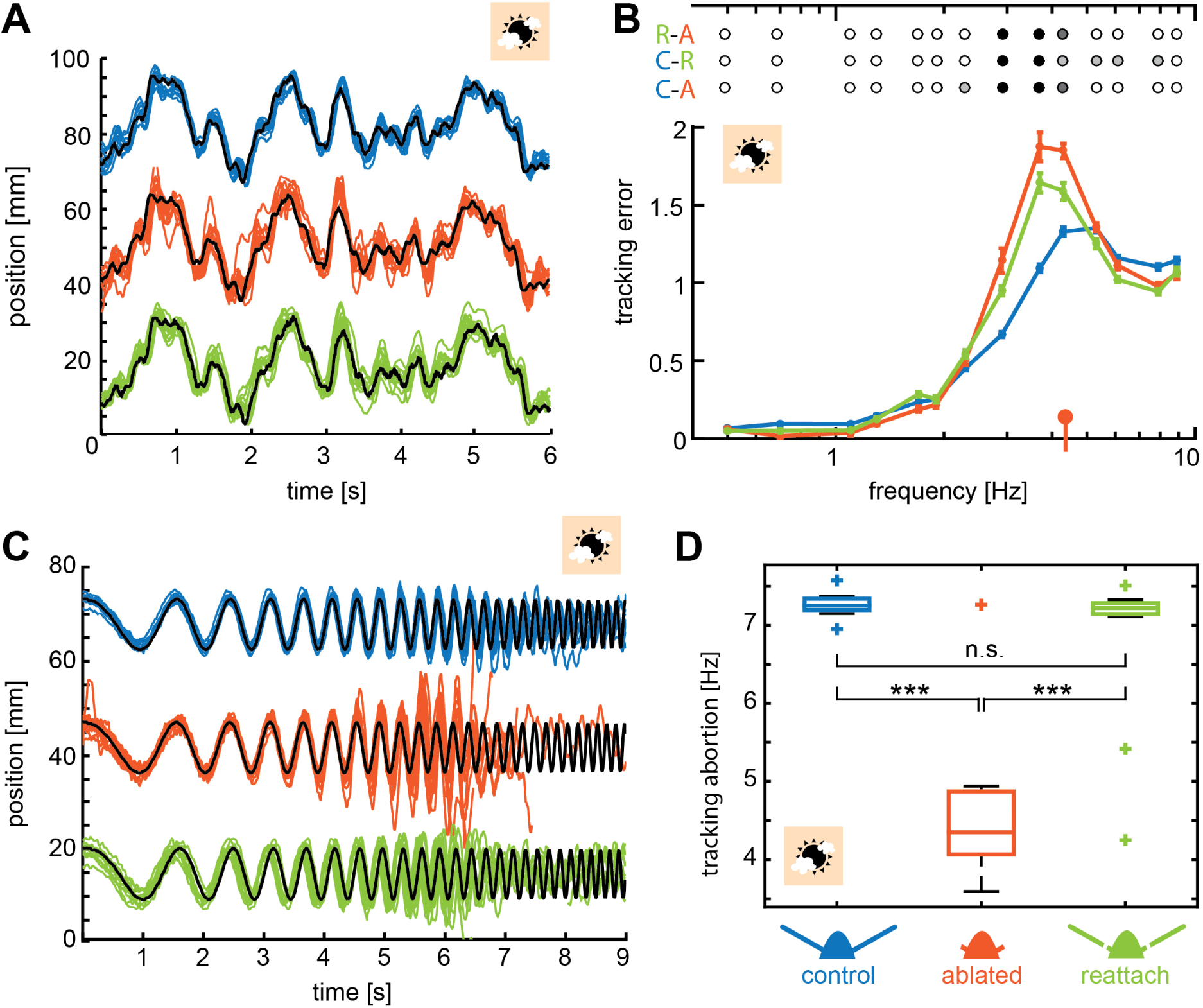
Flagella ablated hawkmoths showed reduced tracking performance of flowers moving at high frequencies. **A, C**. Trajectories of hawkmoths tracking moving flowers with the sum-of-sines (**A**) and the chirp (**C**) stimulus. Trajectories of the different antennal conditions are stacked for comparability. When tracking a moving robotic flower at 3000 lux, hawkmoths with ablated flagella often overshot the movements of the flower, specifically at higher frequencies. With increasing frequencies, moths also increasingly lagged behind the phase of flower movements more strongly. While the amplitude in the sum-of-sines stimulus was adjusted such that moths of all conditions could track the entirety of the stimulus, the chirp stimulus forced moths with too large overshoots and phase-lags to loose contact with the flower, and abort tracking (see red tracks in **C**). **B**. Together, overshooting and phase-lags resulted in an increased tracking error of flagella ablated moths following the sum-of-sines stimulus at frequencies between 2 and 6 Hz, compared to both the control and re-attached condition. Linear mixed-effects models were used to compare the tracking error of the different antennal treatments with respect to frequency, colours indicate significance (black p<0.001, dark grey: p<0.01, grey p<0.05, white p>0.05, Table S3.1). The red indicator on the x-axis gives the median frequency at which flagella ablated moths aborted tracking the chirp-stimulus (**D**). Curves show the mean and 95% confidence intervals of the mean, calculated in the complex plane. **D**. For the chirp stimulus, we compared the movement frequency of the flower, at which the moths aborted tracking, across antennal treatments, showing that flagella ablated moths lost contact with the flower at significantly lower frequencies than the control and re-attached condition. A Friedman test was used to compare between the treatments (*** p<0.001, **: p<0.01, * p<0.05, Table S3.2). See also Fig. S3.1, S3.2

Tracking performance of hawkmoths in the control condition was consistent with the conclusions of previous investigations of intact individuals of this species [Farina et al. 1995, Farina et al. 1994, Stöckl et al. 2017]. In all antennal conditions, the control moths tracked the sum-of-sines stimulus accurately at low frequencies (Fig. 3A): at 3000 lux, their tracking errors were close to 0 for flower movements up to 1 Hz, indicating nearly perfect tracking (Fig. 3B). With increasing frequency, tracking errors increased, but there was no significant difference in tracking error between antennal conditions for frequencies below 2 Hz (Fig. 3B, Table S3.1). At higher frequencies, flagella ablated moths overshot the flower movements, resulting in a greater lag between the position of the moth and the flower and thus larger tracking errors (Fig. 3B): in the frequency range of 2 to 5 Hz, tracking errors were significantly higher for the flagella ablated moths than for both the control moths and moths with re-attached flagella (Table S3.1). In this frequency range, hawkmoths with re-attached flagella also had significantly higher tracking errors than control moths. Thus, the reduction of antennal mechanosensory feedback impaired flight control specifically at the higher temporal frequencies of flower movement, which compel the moths to perform fast turns. The ability of flagella ablated moths to track at frequencies below 2 Hz suggests that vision (and possibly other sensory modalities) provide feedback that is sufficiently fast to enable control of slower manoeuvres.

To ensure that the differences in flight performance between the three antennal conditions was independent of the specific type of flower movement, we presented the moths with a “chirp” stimulus in which the amplitude of flower movement was held constant at 20 mm, while its frequency continuously increased from 0 to 7.3 Hz (Fig. 1D). At low frequencies, hawkmoths of all antennal conditions tracked this stimulus with high fidelity, as indicated by the low tracking error (Fig. 3C). However, as the flower frequency increased, flagella ablated moths tended to overshoot the position of the flower at the end of each sideways movement, when the flower movement changed direction. The accumulated phase lag and overshoot were eventually large enough to cause the moths to lose contact with the nectary, and abort flower tracking (Fig. 3C). Only 1 out of 12 ablated moths succeeded in following the flower movement during the entire stimulus at 3000 lux. In contrast, all control and 10 out of 12 re-attached moths tracked the flower until the maximum frequency. As a measure of flight performance, we quantified the frequency at which hawkmoths in the different antennal conditions aborted flower tracking (Fig. 3D). At 3000 lx, there was no significant difference between the control and re-attached flagella moths (Table S3.2). Flagella ablated hawkmoths aborted flower tracking at a median frequency of only 4.4 Hz, significantly lower than the other two conditions (Fig. 3D, Table S3.2).

Despite the differences between the movement patterns and their demands on flower tracking, we observed similar trends in hawkmoth flight performance for both stimuli. Moths in all antennal conditions tracked flower movements well at low frequencies; whereas flagella ablated hawkmoths were significantly impaired at higher frequencies compared to the other two antennal conditions. The average frequency at which ablated moths failed to track the chirp stimulus was consistent with the frequency range for which the tracking error with the sum-of-sines-stimulus was greatest, despite the difference in flower velocity between stimuli. These data show that antennal feedback is crucial for fast turns - or directional changes - which are associated with changes in body posture.

### Slow visual feedback impaired hovering and tracking performance of all antennal conditions

In the experiments described so far, hawkmoths of all antennal conditions did not differ in their flight performance at lower frequencies (below 0.7 Hz for stationary hovering and below 2 Hz for flower tracking). At these frequencies, feedback from other sensory modalities likely mitigates the problems in flight control caused by antennal ablation. In particular, visual feedback is known to provide information about changes in insect body (head) position in flight (for a review see [Srinivasan et al. 1999]), albeit with longer latencies and a lower frequency range than mechanosensory feedback [Sane et al. 2007]. Because the latency of visual feedback depends on the ambient light intensity [Stöckl et al. 2017], we next tested how the reliability of visual feedback affects the hawkmoth’s flight performance by changing the ambient light intensity, in combination with the antennal manipulations. Specifically, we tested the same group of hawkmoths at an illumination of 30 lux, close to the light intensity limit at which these diurnal hawkmoths are still able to reliably approach and feed from the artificial flowers [Stöckl et al. 2017]. If flagella ablated hawkmoths relied mainly on visual feedback for flight control when they lack antennal mechanosensory feedback, their flight performance should be poorer under low light, as compared to bright light. However, we did not expect any difference in the performance of hawkmoths in the control and re-attached conditions, because these moths receive fast feedback from their antennal mechanosensors.

As expected, at lower frequencies the tracking error of flagella ablated hawkmoths subjected to the sum-of-sines stimulus was greater in dim light than in bright light (Fig. S3.1B). However, we observed the same effect in the control and flagella re-attached conditions (Fig. S3.1A, C). Experiments using the chirp stimulus further confirmed that the low light intensity affected tracking performance in all antennal conditions (compare Fig. 3D and Fig. S3.2). To compare the performance of moths in dim and bright light, we measured the difference in the frequency at which each moth reached a tracking error of 1 with the sum-of-sines stimulus (Fig. 4A), as well as the difference in the frequency at which moths aborted tracking with the chirp stimulus (Fig. 4B). For both stimuli, we did not find significant differences between antennal conditions (Table S4.1 & S4.2), indicating that visual feedback did not compensate for the loss of mechanosensory feedback in flagella ablated moths. Instead, the slower visual processing affects flight control simiarly in all antennal conditions.

**Fig. 4.**
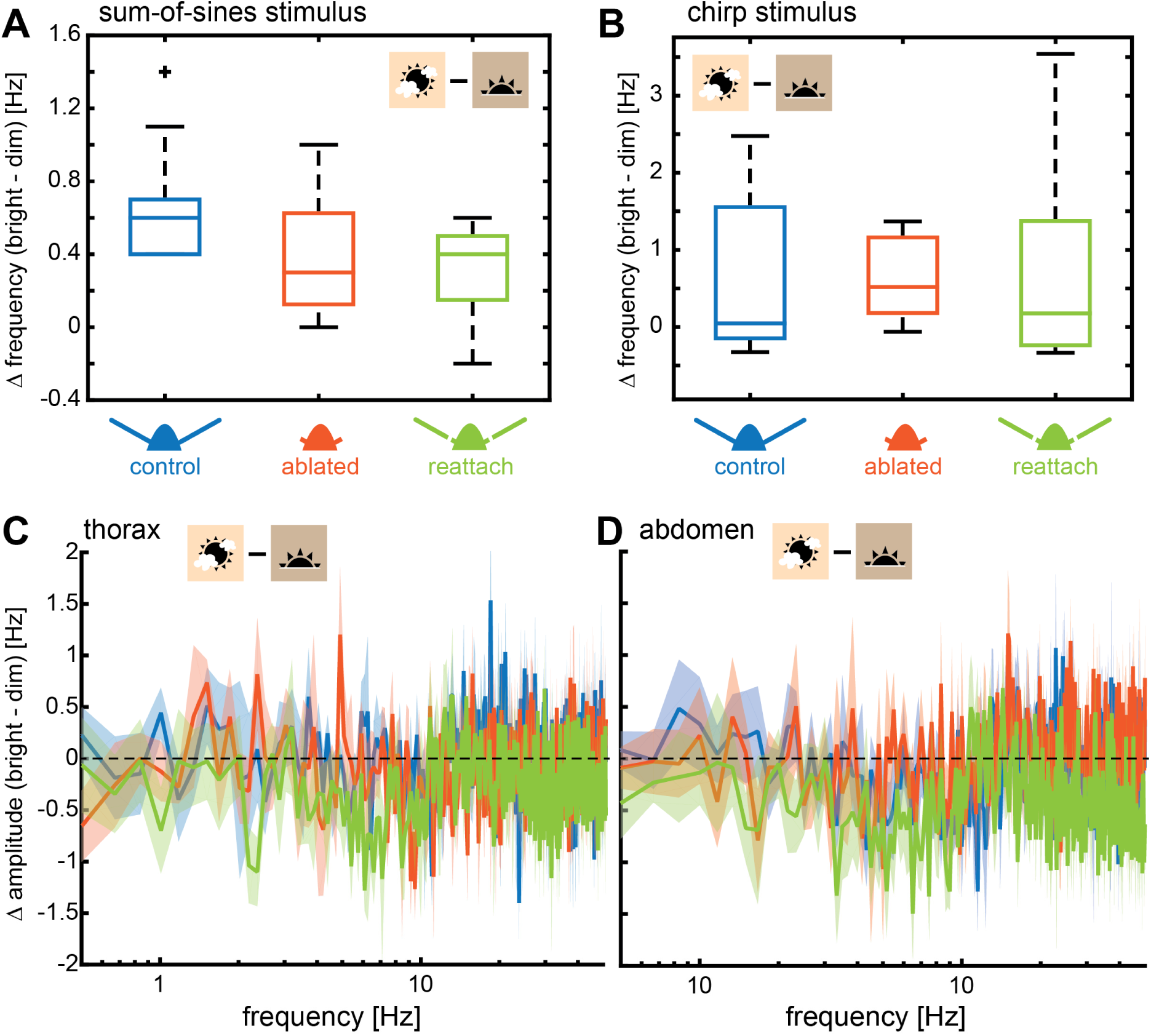
Light intensity had the same effect on all antennal treatments. To test the effect of visual feedback, and its possible interaction with antennal mechanosensory feedback, on flower tracking, we performed all experiments both in bright (3000 lux) and dim (30 lux) light intensities. Hawkmoths showed reduced tracking performance of artificial flowers moving at higher frequencies in dim light, due to the slowing of the visual system (Fig. S2.2, S3.1., S3.2). Here we compare tracking performance between bright and dim light across antennal treatments. **A**. We quantified the difference in frequency between light intensities at which moths reached a tracking error of 1 with the sum-of-sines stimulus. There was no significant difference (Table S4.1) between antennal conditions, suggesting that vision reduced tracking performance in dim light irrespective of the presence or absence of mechano-sensory feedback. **B**. Similarly, there was no significant difference between the tracking performance in dim and bright light for the chirp stimulus (quantified as the difference of tracking abortion frequency at the two light conditions) (Table S4.2). **C-D**. We determined the difference between the log-transformed magnitude spectra for thorax (**C**) and abdomen (**D**) jitter in bright and dim light. No significant effect of antennal condition was found using Friedman comparisons of the average difference in thorax or abdomen movements (Tables S4.3 & S4.4).

This hypothesis was further supported when we compared the moths that were stationary hovering in bright (Fig. 2) and dim light (Fig. S2.2). To quantify the differences in flight performance between the two light intensities, we calculated the difference in the amplitude of thoracic and abdominal jitter across a range of frequencies in dim and bright light for individual moths (Fig. 4 C, D). We found no significant effect of antennal condition on the average difference in thorax (Table S4.3) or abdomen (Table S4.4) jitter between dim and bright light. In conclusion, as in the experiment with the moving flower, the stationary hovering experiment did not reveal any effects of light intensity (causing differences in visual reliability) on flight performance in flagella ablated hawkmoths that we did not observe in control moths as well.

## Discussion

Although visual and mechanosensory feedback is known to play a prominent role in the control of insect flight, it is not clear how these inputs are integrated by the insect brain to generate behaviour. In Dipteran flies, which use halteres as gyroscopic sensors, vision and mechanosensation operate in frequency ranges that are complementary [Mureli & Fox 2015, Yarger & Fox 2016]]. A natural question arising from these studies is: how do insects that lack halteres process mechanosensory and visual feedback? To address this question, we here investigated how visual inputs from compound eyes and mechanosensory inputs from antennal Johnston’s organs control flight in combination. For both stationary hovering and flight manoeuvres during flower tracking in *Macroglossum stellatarum*, our data show that antennal mechanosensory input is crucial for control of fast flight manoeuvres, while visual input controls the slower ones - similar to observation in flies.

### Flagellar re-attachment restores flight performance

We have shown that flight control in the diurnal hawkmoth *M. stellatarum* requires feedback from antennal mechanosensors. As also observed in previous experiments by [Sane et al. 2007], re-attaching the flagellum restored flight performance by reloading the Johnston’s organs, both for stationary hovering and flower-tracking behaviours. This is consistent with the growing body of evidence [Dieudonné et al. 2014, Gewecke & Niehaus 1981, Niehaus 1981, Sane et al. 2010] that Lepidoptera use antennal Johnston’s organs for flight control.

As demonstrated previously [Sane et al. 2007], while flagellum re-attachment improved flight performance significantly compared to the flagella ablated condition, it did not restore it to the level of intact animals. One reason for this observation might be that some properties of the re-attached flagella differ from those of intact animals. Re-attached flagella are not connected to the haemolymph system of the hawkmoth and thus dry out, which reduces their weight by more than 50% (see Methods). Since the flagella are thought to provide a mass that inertial forces act on [Sane et al. 2007], changes in weight may considerably alter the sensory input to the Johnston’s organs. Changes in the flexibility of the flagella due to moisture loss may also contribute to this effect. Moreover, there are also mechanosensors along the length of the flagellum, which may be important for flight control. When the antennal nerve is severed, these mechanosensory units remain inactive even after flagellar reattachment, which may add to the observed deterioration in their ability to control flight. The roles of these mechanosensors and of the weight and flexibility of the flagella needs to be further explored in future experiments.

### Antennal mechanosensation and vision operate in different frequency bands

Our experiments quantified the frequency range in which antennal mechanosensory feedback is required for the control of flight in *M. stellatarum* moths. We demonstrate that flagella ablated hawkmoths can track flowers moving at frequencies below 2 Hz with the same fidelity as hawkmoths with intact antennae (Fig. 2 & 3). This suggests that control of slower manoeuvres is not as dependent on antennal mechanosensory feedback, as is the control of faster manoeuvres. On the other hand, flagella ablated hawkmoths performed significantly worse than moths with intact and re-attached flagella at flower movements above 2 Hz, where more rapid turns are required to follow the lateral trajectory of the moving flower. Our findings are mirrored in the study of Dipteran flight control: slower rotations of fruit flies are tuned stronger to visual feedback, whereas faster rotations require feedback from haltere mechanosensors [Sherman & Dickinson 2003].

Thus, we conclude that mechanosensory feedback from the antennae is essential for the control of fast flight manoeuvres, which require corrective movements to occur in timescales that may not be sufficient for the transduction of visual feedback. This again is analogous to the finding that the control of fast saccadic rotations in Dipterans mainly requires mechanosensory feedback from the halteres, while vision plays a relatively marginal role [Bender & Dickinson 2006, Sherman & Dickinson 2003].

### Vision does not compensate for the loss of antennal mechanosensation in hawkmoth flight control

Both vision and mechanosensation contribute to insect flight control, and the mechanistic underpinnings of this multimodal integration are subject of many ongoing investigations. In Dipteran flies, vision and haltere mechanosensation operate in complementary frequency ranges, and while both inputs are required for stable flight under most circumstances [Yarger & Fox 2016], they do not seem to compensate for each other [Mureli & Fox 2015]]. Antennal movements also depend on feedback from multiple sensory modalities. For example, in honeybees, airflow on the antennae and optic flow influence antennal positioning in tethered as well as free flight [Khurana & Sane 2016]. In the Oleander hawkmoth *Daphnis nerii*, visual feedback modulates antennal positioning in a similar way [Krishnan & Sane 2014].

Here, we tested how vision and antennal mechanosensation in combination influence flight control during flower tracking. Using a bright and a low light intensity, we manipulated the temporal resolution of visual responses [Stöckl et al. 2017, Stöckl et al. 2016]. In dim light, the low speed and reduced reliability of the visual input to flight control causes larger tracking errors when flowers move at high frequencies, for all antennal conditions (Fig. 4). This effect is explained by the fact that visual input is essential for moths to identify and track the flower movement relative to their own position – antennal mechanosensors cannot provide the required information (Fig. 1A). Because visual processing is slower in dim light, moths face greater difficulties in resolving fast flower movements, which causes failure in tracking [Sponberg et al. 2015, Stöckl et al. 2017]. While recent work by [Roth et al. 2016] have demonstrated that mechanoreceptors on the proboscis can play a role in monitoring flower position, in addition to visual input, these results were obtained in the long-tongued hawkmoth *M. sexta*. Our experiments did not test this possibility, but flight ability was also impaired during flower approach in dim light (Fig. S2.1), during which the proboscis provides no positional cues. Moreover, in the short-tongued *M. stellatarum*, proboscis mechanoreception has been shown to be of minor importance for controlling hovering flight [Goyret & Kelber 2011, Zhou 1991]), suggesting that the effects we observed were mainly due to the slowing of the visual input, with little or no contributions from proboscis mechanoreception.

We did not observe a specific effect of light intensity on flight control in the flagella ablated moths. This suggests that, even at higher resolution under brightly lit conditions, visual feedback is unable to mitigate the instability caused by the loss of antennal mechanosensory feedback. Two main hypotheses could explain this finding: first, the contributions of vision and mechanosensation contribute to the motor outputs via separate parallel pathways, whose functions do not overlap. This is unlikely, as recent recordings of descending neurons in Oleander hawkmoth show that they respond to both visual and mechanosensory stimulation [Mohan et al. 2017]. Alternatively, vision and mechanosensation share descending pathways but operate in different frequency ranges, and the visual input is too slow to compensate for the lack of antennal mechanosensory feedback. The latter hypothesis is consistent with physiological studies showing that mechanosensors in Johnston’s organ respond to antennal displacements at frequencies of up to 100 Hz in the hawkmoth *M. sexta* [Sane et al. 2007], whereas the wide-field motion-sensitive neurons of the same species cease to respond at temporal frequencies above 20 Hz [Stöckl et al. 2017], at which most mechanosensors of the Johnston’s organ only show a weak response. Eventually, an assessment of the physiological responses of descending neurons that activate the flight muscles is required to reveal the mechanisms of integration of visual and mechanosensory information in control of flight in hawkmoths.

## Conclusion

Antennal mechanosensation represents one strategy for flying insects to obtain rapid sensory feedback about changes in self-motion, which is crucial for flight control. We showed here that in the diurnal hawkmoth *M. stellatarum*, mechanosesory feedback from antennae is required for the control of fast light manoeuvres and rapid deviations from their hovering position, whereas their visual system drives the control of slower flight manoeuvres. These findings detail a striking similarity to the interaction between mechanosensory halteres and vision in the Dipteran flight control model, and for the first time dissect the combined role of visual and antennal mechanosensory feeback for flight control in hawkmoths, which may be representative for many other non-Dipteran insects.

## Methods

### Animals

Wild adult *Macroglossum stellatarum* L. (Sphingidae), were caught in Sorède, France. Eggs were collected and the caterpillars raised on their native host plant *Gallium sp*. The eclosed adults were allowed to fly and feed from artificial flowers similar to the experimental flowers, in flight cages (70 cm length, 60 cm width, 50 cm height) in a 14:10 h light:dark cycle for at least one day before experiments.

All animals were tested with intact antennae first (*control*), then with ablated flagella (*ablated*), and finally with re-attached flagella (*reattach*) as described below (Figs. 1, S.1.1). Only data from animals that could be tested under all three antennal conditions was included in the final data analysis.

### Surgery: Flagella ablation and re-attachment

For flagella ablation, moths were held, by their thorax under a dissection microscope and their flagella were clipped with a pair of surgical scissors, while retaining 5-10 annuli (Fig S1.1B). This ensured that Johnston’s organs, located at the base of the antennae, were left intact but unloaded. Ablated flagella were preserved in a plastic petri dish with wet tissue to prevent them from drying and losing shape until they were re-attached to the same individual. Moths were left to recover from the surgery and tested on the following day.

To re-attach the flagella, moths were immobilized by cooling at 3° C for 8 minutes, followed by 2 min at −20° C. Flagella were quickly attached to the flagellar stump with a small amount of superglue (Loctite Super Glue Gel, Henkel, Fig. S1.1C). After ensuring that the flagella were properly attached, moths were placed inside a plastic box (10cm × 10cm × 8cm) on a wet tissue paper for 10 min to keep them quiescent and ensure proper reattachment. In case an animal broke the re-attached flagella, a spare one of similar size was used to repeat the re-attachment procedure. Moths were then allowed to recover for a day, before being used in experiments.

We noticed that the flagella lost moisture once re-attached. To quantify the reduction in weight due to moisture loss, we weighed a set of flagella directly after surgery and a few days later when they had dried. Dry flagella had significantly lower weights than freshly ablated flagella (moist: 1.2 ± 0.2 mg, dry: 0.4 ± 0.3 mg; median and inter-quartile range, Wilcoxon rank sum test, z-value = −5.915, p<0.001). We could not determine the weight of the glue used for reattachment, but it is unlikely to exceed the difference between dried and moist flagella, considering the tiny amount of glue used.

### Experimental setup

We used a robotic flower assay as our experimental setup. This assay was first pioneered by [Farina et al. 1994, Sponberg et al. 2015], also used in [Stöckl et al. 2017]. A flight cage (same size as the holding cage) was lined with soft muslin cloth and covered with black cloth on the outside, on three sides, while the front and top were sealed with Perspex windows to allow filming. An artificial flower (48 mm in diameter, on a 140 mm stalk) at the centre of the flight cage, with a nectary (opening of 8.3 mm diameter) filled with 10% sucrose solution, could be moved sideways (in arcs around the central pole), controlled by a stepper motor (0.9 degree/step resolution, 1/16 microstepping, Phidgets, Inc.) and a custom-written Matlab program. The cage was illuminated from above with an adjustable white LED panel and diffuser (CN-126 LED video light, Neewer). Intensity was set to 3000 lux for the bright light condition and 30 lux for the dim light condition (measured with a Hagner ScreenMaster, B. Hagner AB, Solna, Sweden, at the position of the artificial flower). In addition, two 850 nm IR LED lights (LEDLB-16-IR-F, Larson Electronics) provided illumination for the infrared-sensitive high-speed video cameras (MotionBLITZ EoSens mini, Mikrotron) used to film the flower and moths. Videos were recorded at 100 fps, allowing us to record sequences of up to 28 seconds, which were required for our analysis of flower tracking. One camera was placed on top of the cage to film the flower and moth from above during all tests. For experiments with the stationary flower, a second camera providing a rear view was placed on a tripod outside the experimental cage, at approximately 30 cm distance from the artificial flower.

### Behavioural experiments

Eclosed moths were taken from their holding cage and placed in small individually marked cardboard boxes, in which they would be held between trials. For the duration of the experiment, moths were only given access to sucrose from the artificial flower during trials in the experimental cage. A single hawkmoth at a time was introduced into the experimental cage.

We performed two sets of experiments: one, in which we filmed the moth’s approach to and hovering at a stationary flower with both the top and the rear camera, and one in which we filmed the moth tracking a moving flower using only the top camera. In this second set of experiments, we started moving the artificial flower once the moth began to feed from it. We used two different types of movements, the “sum-of-sines” stimulus and the “chirp” stimulus, in the same flight bout. The first 16 s of the sequence thus comprised of the pseudo-random sum-of-sine stimulus composed of the following 14 frequencies, which were prime multiples of each other to avoid harmonic overlap: 0.5, 0.7, 1.1, 1.3, 1.7, 1.9, 2.3, 2.9, 3.7, 4.3, 5.3, 6.1, 7.9, 8.9 Hz. High frequencies had lower amplitudes and vice-versa, to assure equal velocities at all frequencies (Fig.1D). The sum-of-sines stimulus was followed by a brief stationary phase of 0.5 s, and then the 10 s lasting chirp stimulus with fixed movement amplitude of ≈20 mm and frequencies increasing over time from 0 up to 7.3 Hz (Fig.1D) with the relation: frequency = 1.15^(time-1)^.

The protocols were similar for both sets of experiments. Each individual was tested six times: in three antennal conditions (intact *control, ablated* and with *reattached* flagella), and in two light intensities (3000 and 30 lux). Because *M. stellatarum* were less motivated to fly in dim light, we first tested the moths in dim light, when they were hungriest and had the highest motivation to forage, and in bright light (3000 lux) later the same day. If a moth did not track both the sum-of-sines and the chirp stimulus (or the stationary flower for at least 6 s), we repeated the test the next day, until a full set of data was collected and the experiment moved on to the next condition. This experimental strategy gave flagella ablated (and re-attached) moths a chance to adapt to their altered mechanosensory feedback, and practice flying and tracking the flower on several days before succeeding. Indeed, our observations suggest that hawkmoths learned to adjust their flight to the lack or change of mechanosensory feedback, as the initial flight attempts of many flagella ablated (and to a lesser degree re-attached) moths showed more severe impairments than consecutive attempts. Our final dataset includes only individuals that tracked the flower in all three antennal conditions in both light intensities. We used 6 individuals for the experiment with the stationary flower, and 12 different individuals for the experiment with the moving flower.

### Data Analysis

The positions of the flower and the hawkmoth were digitised from the videos using the DLTdv5 software for Matlab [Hedrick 2008, Dyhr et al. 2013]. In experiments with stationary flowers, both the approach and the stationary hovering were digitised, whereas in experiments with moving flower, only sequences during which the proboscis of a moth was in contact with the nectary were rated as ‘tracking’ and digitized (as in [Sponberg et al. 2015, Stöckl et al. 2017]). In the top view, a point on the flower, and a reliably identifiable point on the pronotum of the moth were used for reference. From the rear view videos, we used the centre of the nectary, the centre of the pronotum and the centre tip of the abdomen (Fig. 3).

#### General behaviour

We characterised the general behaviour of all hawkmoths in the moving flower experiments, including those that did not complete all antennal conditions and light intensities, and thus were not included in any further analysis. We classified their behaviour into three different categories (Table 1): non-flying (animals which would not take off after 5 minutes in the experimental cage), flying (animals which flew but would not feed from the flower) and tracking (animals feeding from and tracking the flower, at least partially). We used multinomial logistic regression (package mlogit v0.2-4: [Croissant 2013]) to model the rates of one of the three behaviours (*non-flying, flying, tracking*, Table 1) as a function of antennal condition and light intensity, including the identity of individual moths as a random factor (Table S1).

#### Stationary flower experiments

To compare the stability of hovering flight between the different antennal conditions and light intensities, we analysed the position of the thorax and abdomen for a 6 s interval of hovering at the flower nectary during feeding (given perfect hovering, the thorax should retain a stable position, because the flower was immobile). We quantified the amplitude of thorax and abdomen movements across different frequencies by Fourier transforming their position over time (Fig. 2B,C). To assess the effect of antennal condition across frequencies, we applied a linear mixed-effects model [Bates et al. 2015] with antennal condition, frequency and their interaction as fixed effects and individual identity as a random effect on the log-transformed magnitudes of body movement. We confirmed that the full model did explain the variance better than reduced versions of the model (likelihood ratio test) before performing post-hoc comparisons using the “lmerTest” package in R [Kuznetsova et al. 2017].

To compare these measures across light intensities, we calculated the difference between the log-transformed magnitude spectra of thorax and abdomen position in bright and dim light for each antennal condition (Fig. 4C,D). We then compared these using general linear models of the same form as above.

#### Sum-of-sines movement

We used system identification analysis [Cowan et al. 2014] to characterise hawkmoth flower tracking performance. This analysis is possible because the sum-of-sines stimulus fulfils the requirement of linearity, i.e. it generates the same flower tracking performance at different amplitudes and phase relationships (see Supplement, [Stöckl et al. 2017]. Hawkmoth flower tracking can be described by two components: gain and phase [Farina et al. 1994, Sponberg et al. 2015, Stöckl et al. 2017]. Gain relates the amplitude of flower movement to hawkmoth movement (1 for perfect tracking), while the phase describes the lead or lag of the hawkmoth with respect to the flower movement (0 for perfect tracking). We used a metric called tracking error ε [Roth et al. 2011, Sponberg et al. 2015], which incorporates effects of both gain and phase to quantify tracking performance of hawkmoths (Fig. 3B). It is calculated as the complex distance between the moth’s response *H(s*) and the ideal tracking conditions (gain=1, phase lag=0), where *s* is the Laplace frequency variable:

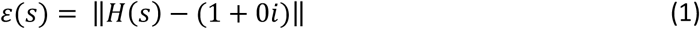

A tracking error of 0 means perfect tracking (comprising a gain of 1 and a phase lag of 0), while the tracking error is 1 if the hawkmoth and flower movement are uncorrelated (e.g. when either the hawkmoth remains stationary and the flower moves, or vice versa). We calculated average tracking errors and their confidence intervals within antennal conditions by averaging data in the complex plane, to avoid artefacts resulting from separating gain and phase components when transforming them and averaging in the non-complex plane (see [Stöckl et al. 2017] for discussion).

Since our tracking error metric is a complex value, and was only transformed into the non-complex plane after averaging across individuals, it is not straightforward to find appropriate statistical tests to compare tracking error (as well as gain and phase) across antennal conditions and light intensities. Linear mixed effects models would be well suited, but complex data might not fulfil all of the assumptions these models are based on. Lacking an alternative, we had to rely on these tests, as did previous studies with the same approach [Roth et al. 2016, Sponberg et al. 2015] to compare the effect of antennal conditions across frequencies as fixed effects, including individual identity as a random effect. We are confident that overall trends identified as significant by these models are indicative of biologically relevant effects, but advice caution when interpreting differences in significance at individual movement frequencies isolated from the overall trend.

To compare tracking performance across light intensities, we calculated the difference in the frequency at which tracking error reached 1 for both dim and bright light intensity within antennal conditions (similar to [Stöckl et al. 2017]), and compared these across antennal conditions (Fig.4A). For statistical comparisons, we used the Friedman test, which is a non-parametric test that accounts for repeated measures.

#### Chirp movement

The chirp stimulus does not fulfil the linearity criterion, because it does not generate the same flower tracking performance at different amplitudes and phase relationships, but rather contains a saturation non-linearity. Hence, the system identification analysis we used for the sum-of-sines stimulus could not be applied [Roth et al. 2011]. We therefore determined the frequency, at which each individual lost proboscis contact with the flower (i.e. failed at tracking the flower) as a measure of flower tracking performance across frequencies. This measure gave an absolute cut-off frequency at which moths could no longer track the oscillating flower. Because this data was non-parametric and included repeated measures, we used a Friedman test to compare the paired data.

To compare the tracking performance across light intensities, we calculated the difference in termination frequency between dim and bright light for each antennal condition (taking the difference for each individual). These differences between light conditions were then compared across antennal conditions using a Friedman test to retain information about the paired data (Fig.4B).

## Acknowledgements

We thank Merry and Leigh Foster for help with capturing the parental moths in France, Michael Pfaff and Joaquin Goyret for performing pilot experiments, Simon Sponberg for installing our robotic flower set-up and for critical comments on the manuscript, Marie Dacke for allowing us to use two high speed cameras, David O’Carroll for inspiring discussions, Karin Nordström for valuable comments on the manuscript, and Eric Warrant for financial support of AS.

## Author contributions

AS, SPS and AK conceived the study, AD performed the experiments, AD digitized the data, JF conceived the statistical analysis, AS analysed the data with input from AD, AK and JF, AS wrote the initial draft of the manuscript, and all authors discussed and commented on the manuscript.

## Supplementary Figures

**Fig. S1.1:**
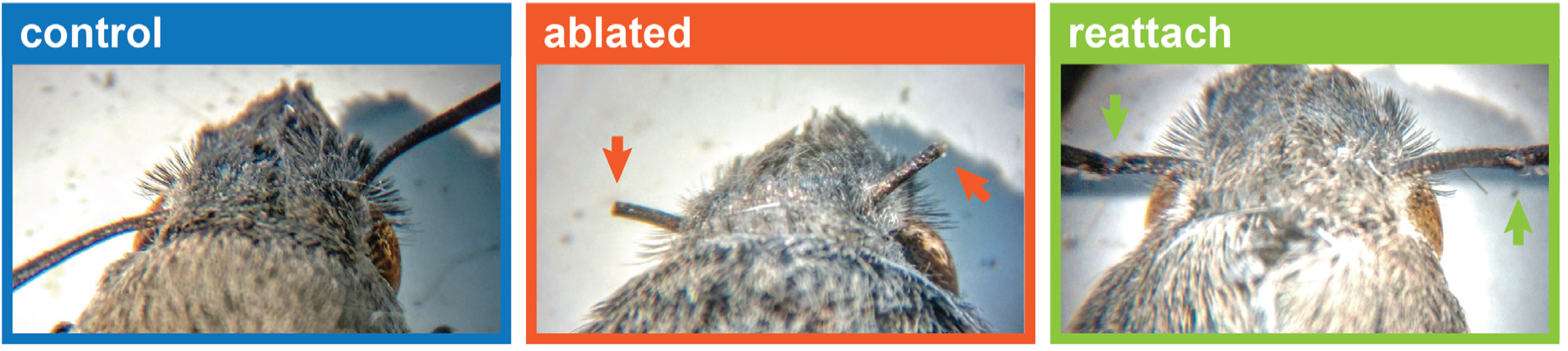
Antennal surgery. To obtain flagella ablated animals, the flagella were cut between the 5^th^ and 10^th^ annulus (red arrows in middle panel). The flagella were preserved and re-attached to the same individual with a small amount of super glue (green arrows, right panel).

**Fig. S2.1:**
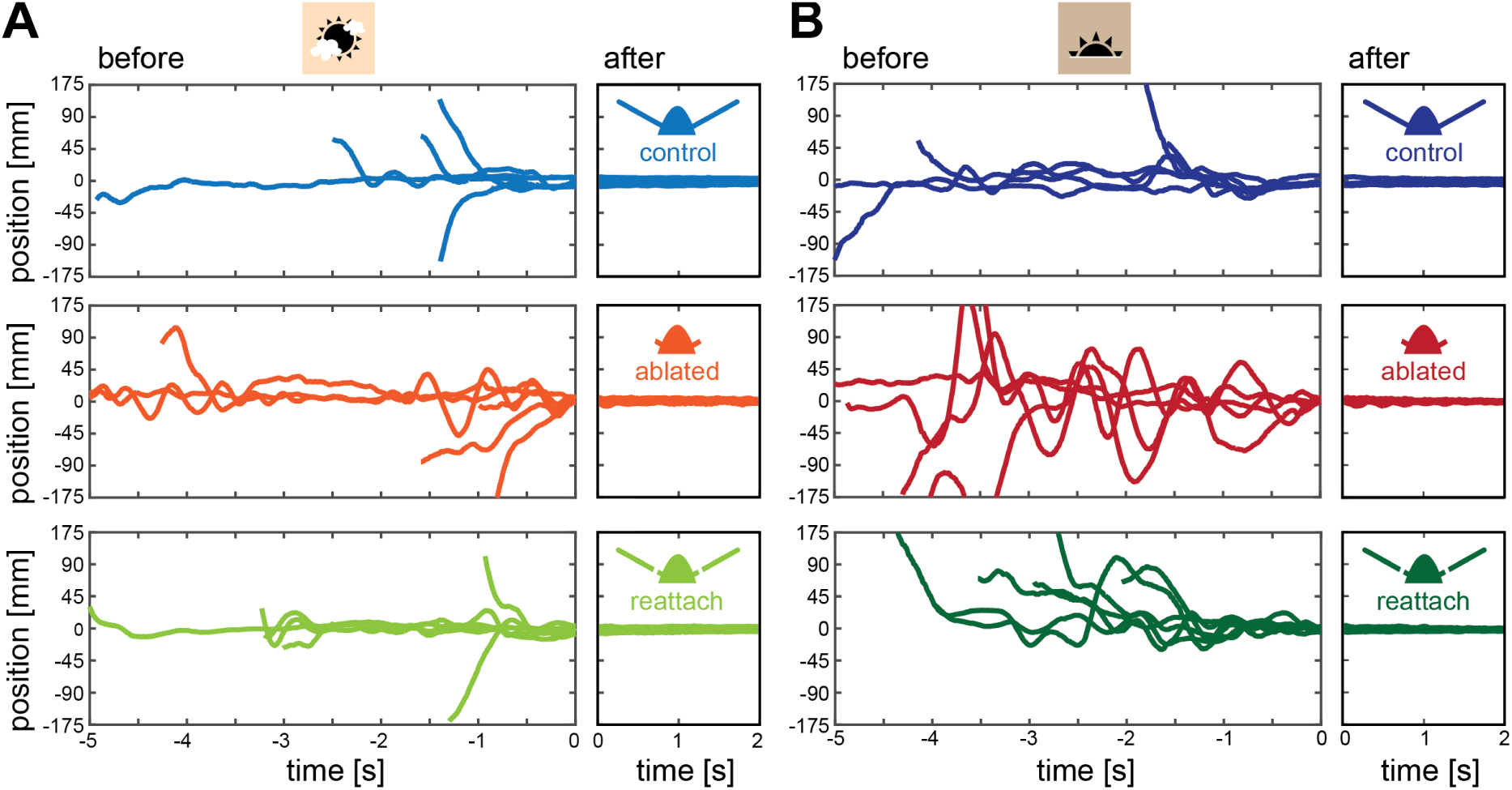
Hawkmoths approaching the moving flower in bright (3000 lux) and dim (30 lux) light. *Before* refers to time intervals before their proboscis made contact with the nectary of the flower, *after* to the period after contact was initiated.

**Fig. S2.2.**
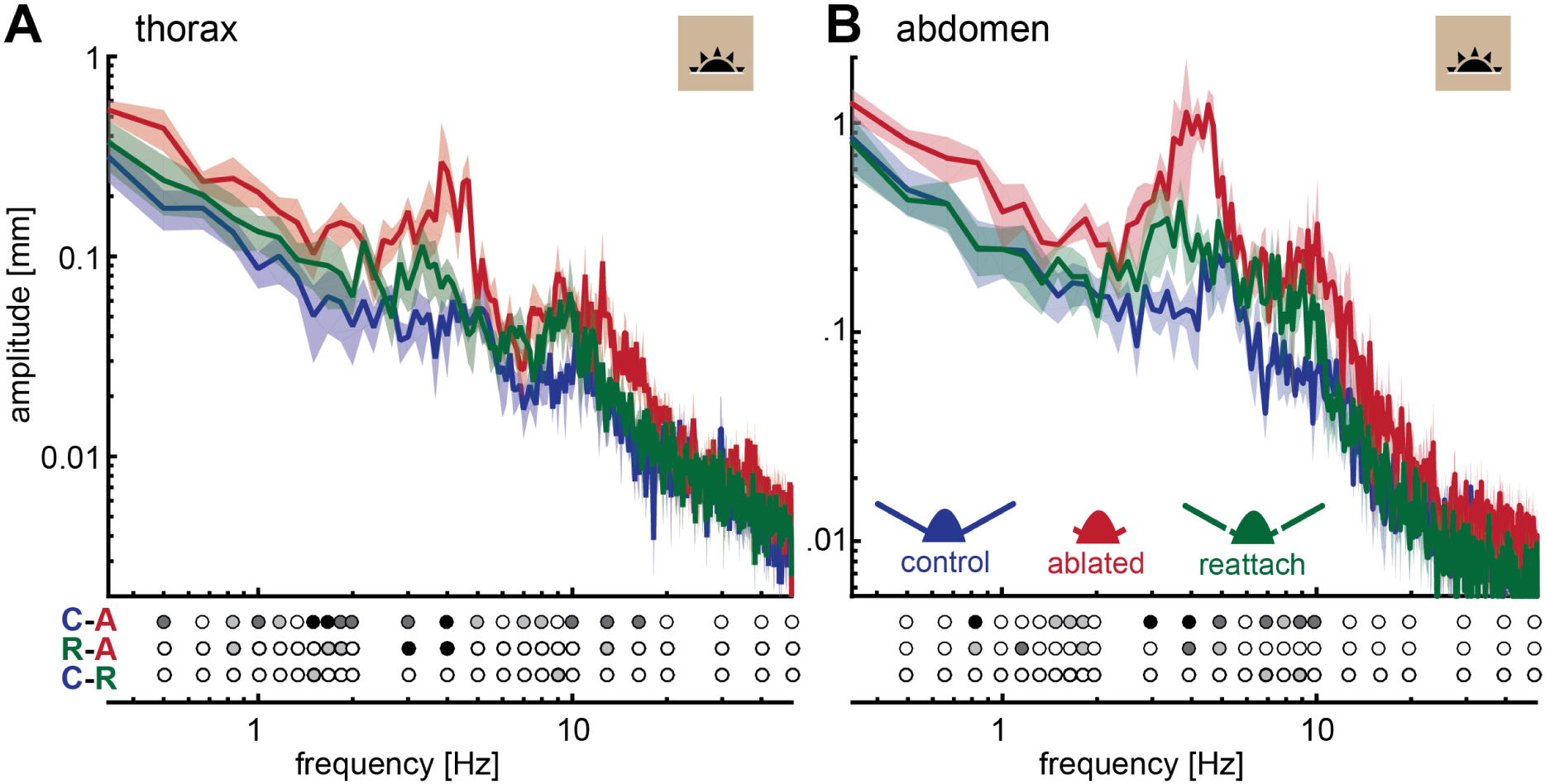
Flagella ablated hawkmoths perform less stable hovering at a stationary flower at 30 lux. Thorax (**A**) and abdomen (**B**) position of moths hovering in front of a stationary flower at 30 lux. Post-hoc tests were performed as part of a general linear model including antennal treatment and frequency (binned to the logarithmic scale) as factors. Statistical significance is indicated below the plots as: black p<0.001, dark grey: p<0.01, grey p<0.05, white p>0.05, see Tables S2.3 and S2.4.

**Fig. S3.1.**
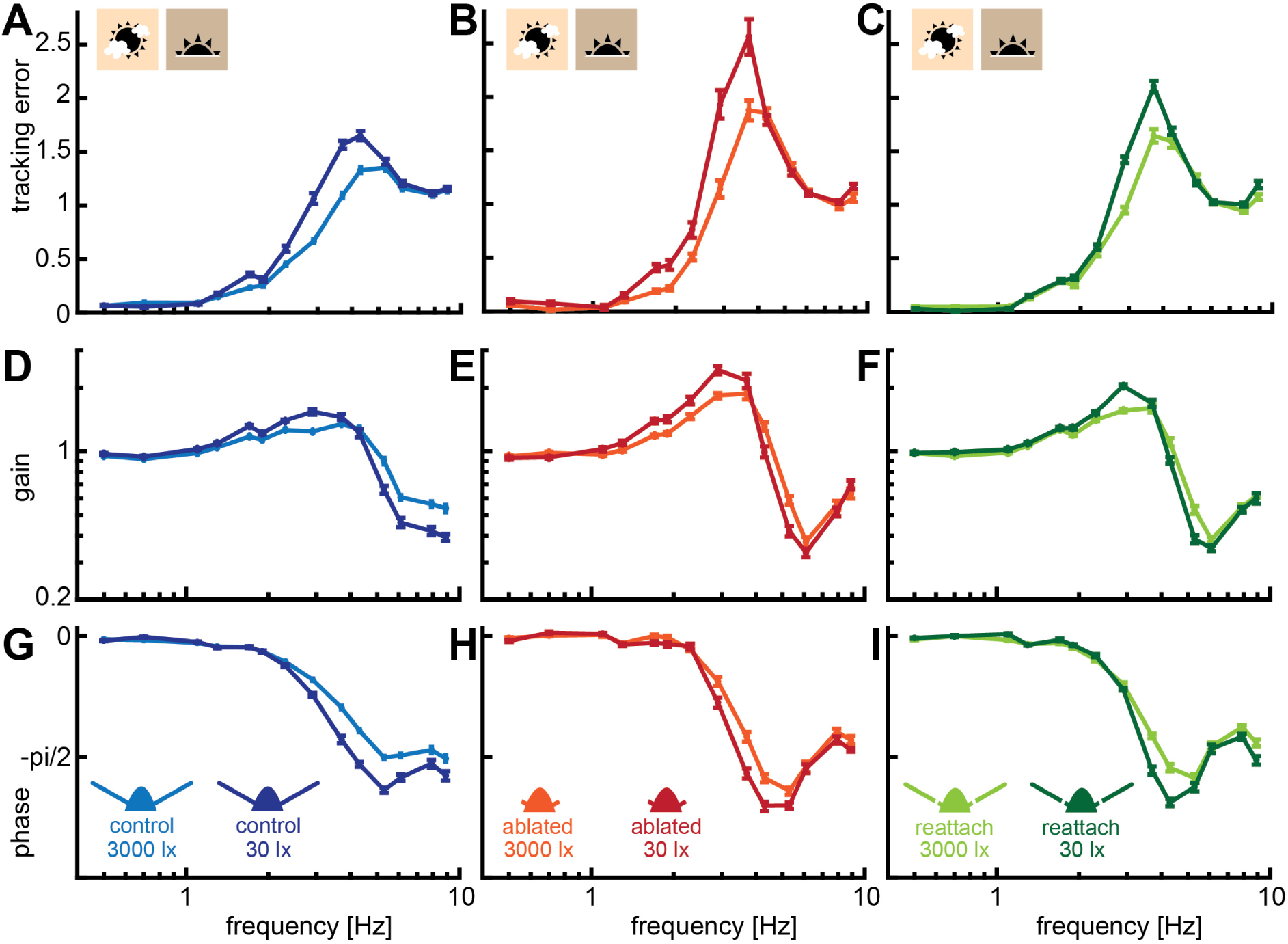
Hawkmoth tracking performance with the sum-of-sines stimulus in bright and dim light. Tracking error (**A-C**), gain (**D-F**) and phase (**G-I**) of hawkmoths tracking a moving robotic flower which moved as a sum-of-sines (see Methods) at 3000 and 30 lux. Curves show the mean and 95% confidence intervals of the mean, calculated in the complex plane.

**Fig. S3.2.**
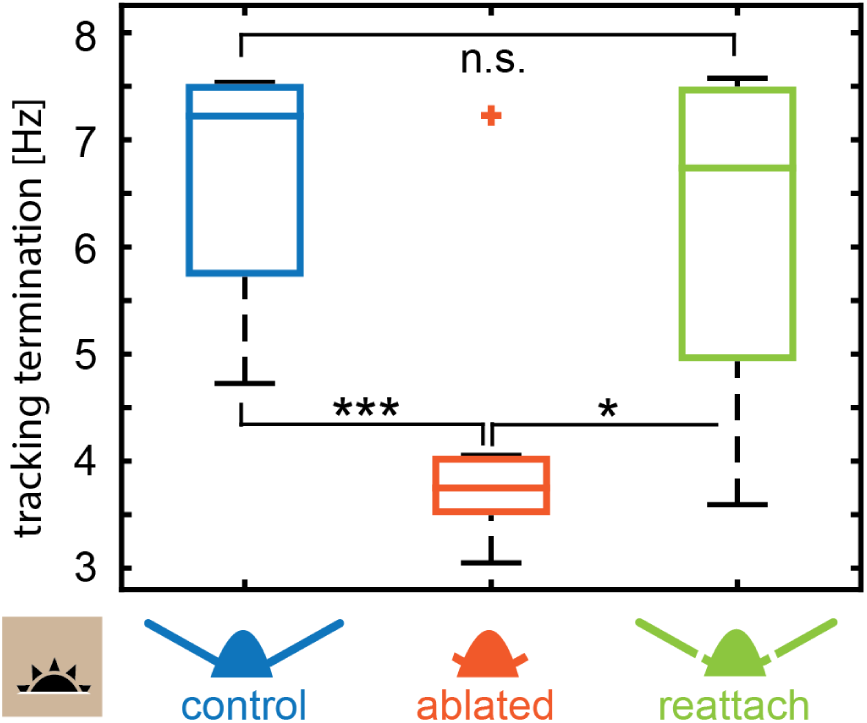
Flower tracking performance of chirp stimuli in dim light. Comparison of the temporal frequency of the flower at which the moths aborted tracking across antennal treatments at 30 lux. A Friedman test was used to compare between the treatments (*** p<0.001, **: p<0.01, * p<0.05, Table S3.2).

## Supplementary Tables

Table S1 Results of the statistical models assessing the effect of antennal treatment and light intensity on the proportion of different behaviours in the flight cages.

The behaviour of each animal was classified into the following categories: *no flight, flight* (but no tracking of the flower), and *tracking*. Some moths were tested multiple times to collect the necessary tracking data, and thus have contributed multiple trials to this dataset. Statistical comparisons were performed using multinomial regression including the identity of individual moths as a random factor, to model the rates of one of the three behaviours as a function of antennal condition and lighting. As no significant interaction between antennal condition and light intensity was found, the fixed effects of the fitted model took the form: behavioural category (no flight, flight, tracking) ~ antennal condition + light intensity. All statistical results are expressed in relation to the probability of observing the *no flight* behaviour in the control condition in bright light.

**Table S1.** Results of the statistical model assessing the effect of antennal condition and light intensity (without interaction) on the proportions of observed behaviours (Table 1).

| Conditions | Coefficient Estimate | t-value | p-value |
| --- | --- | --- | --- |
| *flight* **ablate** - **control** | -1.27 | -1.53 | 0.126 |
| *tracking* **ablate** - **control** | -2.76 | -3.63 | <0.001 |
| *flight* **reatt** - **control** | -1.16 | -1.05 | 0.292 |
| *tracking* **reatt** - **control** | -0.83 | -0.88 | 0.380 |
| *flight* **dim** - **bright** | -1.01 | -1.93 | 0.054 |
| *tracking* **dim** - **bright** | -1.13 | -2.33 | 0.020 |

**Table S2 Results of the statistical models assessing the effect of antennal treatment in the stationary experiment in bright light.** A general linear model was constructed with antennal treatment and frequency (binned to the logarithmic scale) as factors: log(response) ~ antennal condition * frequency + 1|individual

**Table S2.1.** Results of the statistical model assessing the effect of antennal treatment on thorax jitter in bright light (Fig. 2B).

| Treatment | Estimate | t-value | DF | p-value |
| --- | --- | --- | --- | --- |
| control - ablate | -0.664 | -10.92 | 385 | <0.001 |
| control - reatt | -0.002 | -0.04 | 385 | 0.97 |
| ablate - reatt | 0.662 | 10.88 | 385 | <0.001 |

**Table S2.2.** Results of the statistical model assessing the effect of antennal treatment on abdomen jitter in bright light (Fig. 2C).

| Treatment | Estimate | t-value | DF | p-value |
| --- | --- | --- | --- | --- |
| control - ablate | -0.658 | -12.07 | 385 | <0.001 |
| control - reatt | 0.149 | 2.74 | 385 | 0.007 |
| ablate - reatt | 0.807 | 14.81 | 385 | <0.001 |

**Table S2.3.** Results of the statistical model assessing the effect of antennal treatment on thorax jitter in dim light (Fig. S2.2A).

| Treatment | Estimate | t-value | DF | p-value |
| --- | --- | --- | --- | --- |
| control - ablate | -0.742 | -11.63 | 385 | <0.001 |
| control - reatt | -0.257 | -4.03 | 385 | 0.0001 |
| ablate - reatt | 0.485 | 7.60 | 385 | <0.001 |

**Table S2.4.** Results of the statistical model assessing the effect of antennal treatment on abdomen jitter in dim light (Fig. S2.2B).

| Treatment | Estimate | t-value | DF | p-value |
| --- | --- | --- | --- | --- |
| control - ablate | -0.656 | -10.37 | 385 | <0.001 |
| control - reatt | -0.234 | -3.71 | 385 | 0.0002 |
| ablate - reatt | 0.421 | 6.67 | 385 | <0.001 |

**Table S3 Results of the statistical model assessing the effect of antennal treatment on flower tracking performance in moving flower experiments.**

**Table S3.1.** Results of the statistical model assessing the effect of antennal treatment on flower tracking performance with the sum-of-sines stimulus in bright light (Fig. 3B): a general linear model was constructed with antennal treatment and frequency (binned to the logarithmic scale) as factors: log(response) ~ antennal condition * frequency + 1|individual

| Treatment | Estimate | t-value | DF | p-value |
| --- | --- | --- | --- | --- |
| control - ablate | -0.160 | -9.04 | 913 | <0.001 |
| control - reatt | -0.043 | -2.42 | 913 | 0.016 |
| ablate - reatt | 0.117 | 6.62 | 913 | <0.001 |

**Table S3.2.** Results of the statistical model assessing the effect of antennal treatment on flower tracking performance with the chirp stimulus in bright light (Fig. 3D): a Friedman test was performed, with a Tukey-Kramer post-hoc comparison correction for multiple comparisons.

| Treatment | X<sup>2</sup> | p-value |
| --- | --- | --- |
| overall | 19.48 | <0.001 |
| control - ablate |  | <0.001 |
| control - reatt |  | 0.68 |
| ablate - reatt |  | 0.002 |

**Table S3.3.** Results of the statistical model assessing the effect of antennal treatment on flower tracking performance with the sum-of-sines stimulus in dim light (Fig. S3.1): a general linear model was constructed with antennal treatment and frequency (binned to the logarithmic scale) as factors: log(response) ~ antennal condition * frequency + 1|individual

| Treatment | Estimate | t-value | DF | p-value |
| --- | --- | --- | --- | --- |
| control - ablate | 0.184 | -6.36 | 462 | <0.001 |
| control - reatt | 0.028 | 0.98 | 462 | 0.328 |
| ablate - reatt | 0.156 | 5.38 | 462 | <0.001 |

**Table S3.4.** Results of the statistical model assessing the effect of antennal treatment on flower tracking performance with the chirp stimulus in dim light (Fig. S3.2): a Friedman test was performed, with a Tukey-Kramer post-hoc comparison correction for multiple comparisons.

| Treatment | X<sup>2</sup> | p-value |
| --- | --- | --- |
| overall | 16.7 | <0.001 |
| control - ablate |  | <0.001 |
| control - reatt |  | 0.56 |
| ablate - reatt |  | 0.012 |

**Table S4 Results of the statistical model comparing the effect of antennal condition between light conditions for all experiments.** A Friedman test was performed, with a Tukey-Kramer post-hoc comparison correction for multiple comparisons.

**Table S4.1.** Results of the statistical model assessing the effect of antennal condition on the difference in flower tracking error between light conditions with the sum-of-sines stimulus (Fig. 4A).

| Treatment | X<sup>2</sup> | p-value |
| --- | --- | --- |
| overall | 2.71 | 0.258 |
| control - ablate |  | 0.303 |
| control - reatt |  | 0.357 |
| ablate - reatt |  | 0.994 |

**Table S4.2.**
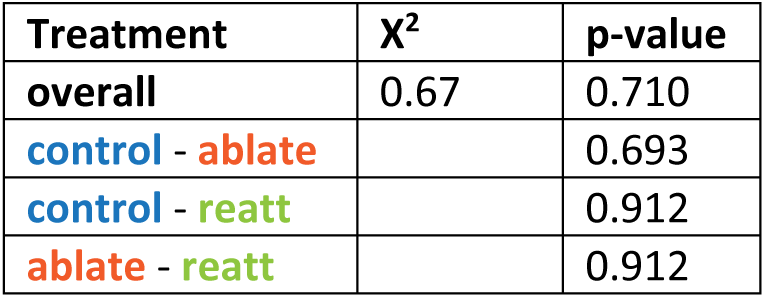
Results of the statistical model assessing the effect of antennal treatment on the difference in flower tracking performance between light conditions with the chirp stimulus (Fig. 4B).

| Treatment | X<sup>2</sup> | p-value |
| --- | --- | --- |
| overall | 0.67 | 0.710 |
| control - ablate |  | 0.693 |
| control - reatt |  | 0.912 |
| ablate - reatt |  | 0.912 |

**Table S4.3.** Results of the statistical model assessing the effect of antennal treatment on the difference in thorax stability during hovering between light conditions (Fig. 4C).

| Treatment | $\chi^2$ | p-value |
| --- | --- | --- |
| overall | 2.33 | 0.311 |
| control - ablate |  | 0.480 |
| control - reatt |  | 0.319 |
| ablate - reatt |  | 0.955 |

**Table S4.4.**
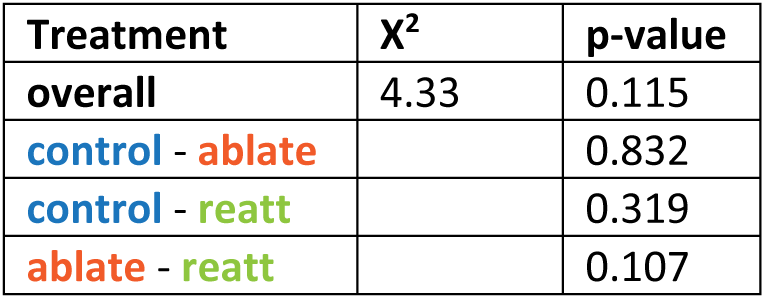
Results of the statistical model assessing the effect of antennal treatment on the difference in abdomen stability during hovering between light conditions (Fig. 4D).

